# Positional effects revealed in Illumina Methylation Array and the impact on analysis

**DOI:** 10.1101/153858

**Authors:** Chuan Jiao, Chunling Zhang, Rujia Dai, Yan Xia, Kangli Wang, Gina Giase, Chao Chen, Chunyu Liu

**Affiliations:** State Key Laboratory of Medical Genetics, Central South University, Changsha, Hunan, 410012, China; Center for Research Informatics, University of Chicago, Chicago, IL, 60607, USA; Department of Psychiatry, University of Illinois at Chicago, Chicago, IL, 60607, USA

**Keywords:** positional effects, DNA methylation, Illlumina Infinium HumanMethylation BeadChips, *ComBat*, batch effects, Infinium Methylation 450K, Infinium Methylation 27K, epigenetics

## Abstract

With the evolution of rapid epigenetic research, Illumina Infinium HumanMethylation BeadChips have been widely used to study DNA methylation. However, in evaluating the accuracy of this method, we found that the commonly used Illumina HumanMethylation BeadChips are substantially affected by positional effects; the DNA sample’s location in a chip affects the measured methylation levels. We analyzed three HumanMethylation450 and three HumanMethylation27 datasets by using four methods to prove the existence of positional effects. Three datasets were analyzed further for technical replicate analysis or differential methylation CpG sites analysis. The pre- and post-correction comparisons indicate that the positional effects could alter the measured methylation values and downstream analysis results. Nevertheless, *ComBat*, linear regression and functional normalization could all be used to minimize such artifact. We recommend performing *ComBat* to correct positional effects followed by the correction of batch effects in data preprocessing as this procedure slightly outperforms the others. In addition, randomizing the sample placement should be a critical laboratory practice for using such experimental platforms. Code for our method is freely available at: https://github.com/ChuanJ/posibatch.

## Introduction

DNA methylation is an important epigenetic modification that regulates gene expression^1^, chromatin structure and stability^2^, and genomic imprinting^3^. DNA methylation has also been implicated in the development of cancer^4-6^ and other diseases^7-9^. Furthermore, several studies indicated that the DNA methylation levels could vary by age^10^, sex^11^, disease affected status^4-9^, circadian rhythms^12^, tissues types^13^ and other factors.

Many methods have been used to measure the methylation levels of cytosines, such as blotting, atomic force spectroscopy, genomic sequencing, bisulfite sequencing, methylation-specific PCR, microarray analysis, etc^14^. The high-throughput methods can be classified into two major categories: next-generation sequencing (NGS) and microarrays. In many NGS-based technologies, the whole-genome bisulfite conversion is generally regarded as a gold standard for highest genomic coverage, accuracy, and resolution. Microarray-based technologies such as Illumina Infinium HumanMethylation27 BeadChip Array (Methyl27)15, Illumina Infinium HumanMethylation450 BeadChip Array (Methyl450)^16, 17^, and Illumina Infinium MethylationEPIC BeadChip microarray^18^, have been widely used for methylome profiling since the first chip came to market in 2006^19^ with its advantages in terms of low cost, modest DNA requirement, and high throughput^20^. However, these methods are unable to interrogate genomic regions outside of the pre-designed probes, thereby limiting the exhaustive screening of the genome.

Methyl450 was one of the most popular and cost effective tools available allowing researchers to interrogate more than 485,000 methylation loci per sample at single-nucleotide resolution^21^. It has twelve sample sections in one array arranged in a six by two format (Fig.S1). While Methyl27 measures the methylation status of over 27,000 CpG sites in the genome using the Type I assay with twelve sample locations arranged by twelve rows (Fig.S1), Methyl450 increased its capacities upon Methyl27 by adding the Type II assay. However, they suffer from errors introduced by probe cross-hybridization^17, 22^, the probe type bias^16^, single nucleotide polymorphisms (SNPs) contaminated probes ^17, 23^ and so on. Filtering out probes with potential errors and adjusting experimental bias have been necessary data pre-processing steps.

There is also ‘positional effects’ that the same sample in different physical positions on the array could be measured as different methylation levels. The earliest mention of the positional effect in the Illumina gene expression microarray analysis did not provide a method for correction except an advisement to randomly set the samples in the array^24^. Since then, a few papers mentioned the possible existence of positional effects by other names such as the ‘Sentrix position effect’, ‘beadchip effect’, ‘slide effects’ or ‘beadchip position on plate effects’, but failed to provide solid evidence about them, nor provide a convincingly effective method to correct the effect^17, 24-26^. Conventional approaches to correct confounders such as the polygenic regression model^27^ have been attempted, but the scientific rationality of the regression model in the randomly distributed effects is problematic^24^. There is also one unsupervised method named Functional normalization (FN) could correct the effect^28^.

In this article, we compared three methods for correcting the positional effects: *ComBat*, linear regression model and functional normalization. *ComBat* adjusts for known batches using an empirical Bayesian method even in small sample sizes, the linear regression model is a classical method to remove known confounders, and FN is an unsupervised method using control probes as surrogates for unwanted variation ^28^.

While the investigation into the positional effects was not thorough, positional effects have rarely been controlled in analysis like batch effects. Controlling batch effects^29-33^ has been a critical practice in data analysis. Illumina HumanMethylation BeadChip platforms have already been implemented in epigenetic studies of cancer and many other diseases with about 883 papers published so far (NCBI GEO database^34^). Few studies had properly addressed the positional effects, which could lead to potential bias particularly when samples were not placed randomly.

In this study, we closely examined the important technical artifact in the Illumina HumanMethylation BeadChip named “positional effects” using multiple datasets of both Methyl27 and Methyl450. We proved the existence and discussed the origin of the effect, the bias it brings to the research result, and the proper solution to adjust this confounder. Specifically, four methodologies were utilized to evaluate the effects, including: identification of CpG sites that are significantly associated with sample position, the relative contribution to overall variation in measured methylation levels, correlation and variation between technical replicates, and significant differential methylation signals between cases and controls. We further tested several methods to control positional effects along with batch effects to ensure that both artifacts can be managed. With that, we are offering a recommendation for the pre-processing of Illumina methylation data.

## Results

### Analysis of Variance (ANOVA) Results of Methylation Levels and Physical Positions

We analyzed the ROSMAP data with 743 samples in 64 arrays (Table 1, Fig. 1, *Material and methods*). After quality control (QC) pre-processing and filtering, 167,384 probes were tested for the correlations between methylation levels and sample physical positions. 14,063 of them were significantly associated with their sample positions by false discovery rate (FDR) q-value < 0.05, while 153,079 loci were associated with batches. After removal of batch effects, the number of CpG loci associated with position increased to 32,144; and the batch-associated sites reduced to zero.

**Figure 1.**
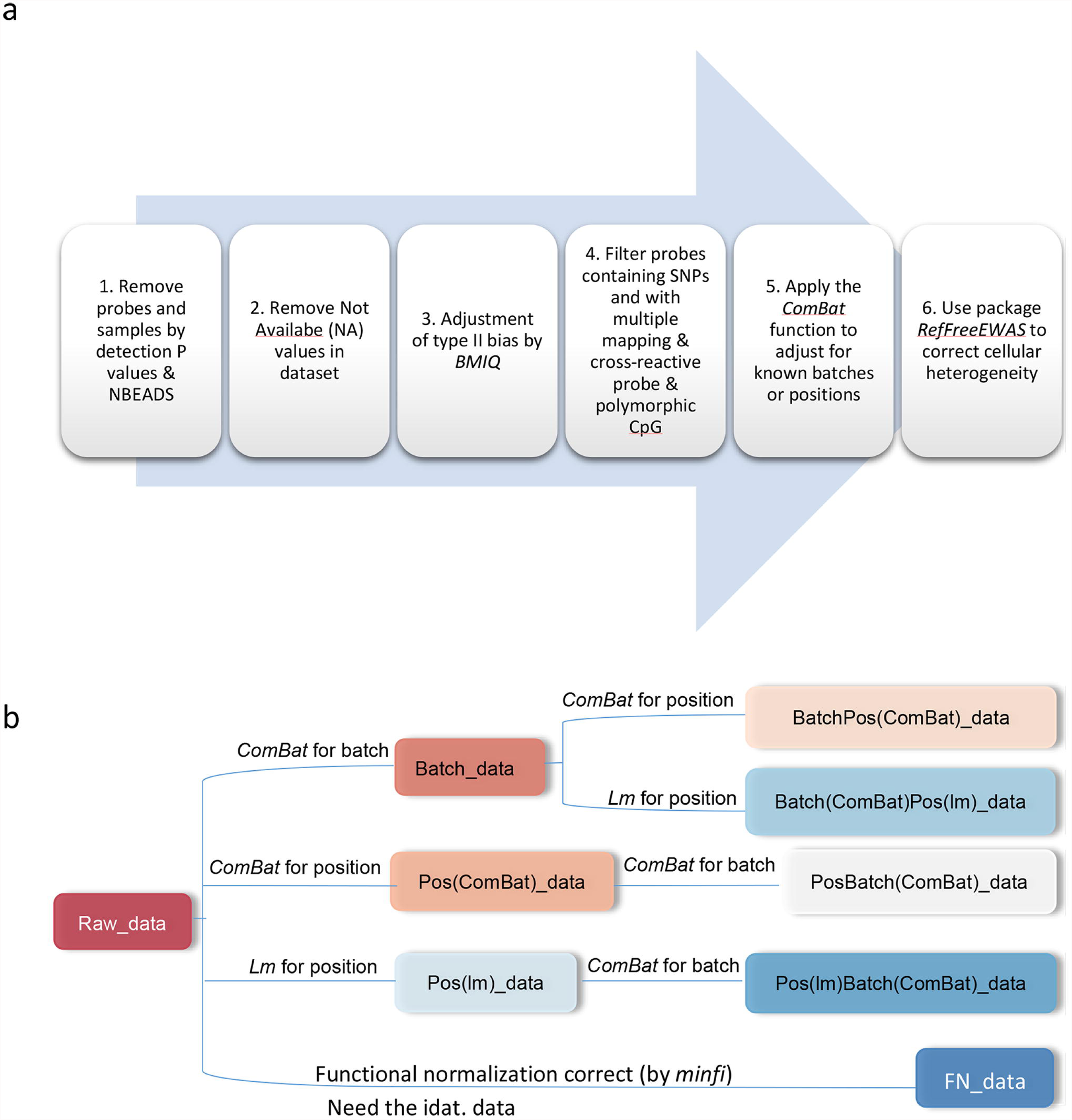
The basic pipeline. (a) The basic pipeline used to process ROSMAP dataset. (b) Datasets in different workflows correcting batch and positional effects. The Fig. 1b is the detailed procedure of Step 5 in Fig. 1a. There are eight datasets in different workflows in Fig 1b, including: Raw_data (data after primary QC and filtering), Batch_data (data corrected the batch effect), Pos(ComBat)_data (data corrected the positional effect by ComBat function), BatchPos(ComBat)_data (data corrected the batch and positional effect sequentially by ComBat in order), PosBatch(ComBat)_data (data corrected the positional and batch effect sequentially by ComBat in order), Pos(lm)_Data (data corrected the positional effect by lm), Batch(ComBat)Pos(lm)_data (data corrected the batch by ComBat and positional effect by lm sequentially, Pos(lm)Batch(ComBat)_data (data corrected the positional effect by lm and batch effect by ComBat sequentially) and FN_data (data corrected by functional normalization by using the preprocessFunnorm function in the minfi package). Note: NBEADS means the number of the beads. BMIQ means Beta Mixture Quantile dilation, a method adjusting the β values of type II probes into a statistical distribution characteristic of type I probes (see Materials and Methods). SNPs means single nucleotide polymorphisms.

**Table 1.** The sample information of ROSMAP data (743 samples). The ROSMAP data is the primary data we used. The samples were from two longitudinal cohort studies at Rush University Medical Center - the Religious Orders Study and the Rush Memory and Aging Project.

| Variable | Number ( or Mean± Standard deviation) |
| --- | --- |
| Sex |  |
| Female | 468 |
| Male | 275 |
| Race |  |
| White | 725 |
| African-American | 14 |
| Native American | 1 |
| Asian or Pacific Island | 3 |
| Age_bl, year | 81.17± 6.98 |
| Age_death, year | 88.01± 6.66 |
| Educational level, year | 16.38± 3.63 |

Abbreviations: age\_bl means age at cycle – baseline which can be the cognitive date, interview date, or clinical evaluation; age\_death means age at death.

We corrected the positional effects only with *ComBat*, and still detected 20 CpG loci associated with positions but left 154,140 probes related to batches. Then the batch and positional effects were sequentially adjusted in two different orders. When corrected for the batch effects first, 21 loci associated with position were identified, and zero associated with the batch. However, when corrected for the positional effects first, 24 position-associated loci were detected, and none of the batch-associated signals were detected.

We noticed that we detected 11,500 CpG loci significantly associated with the batches when we corrected the batch effect first followed by positional effects in BrainCloud dataset (Table 2).

**Table 2.**
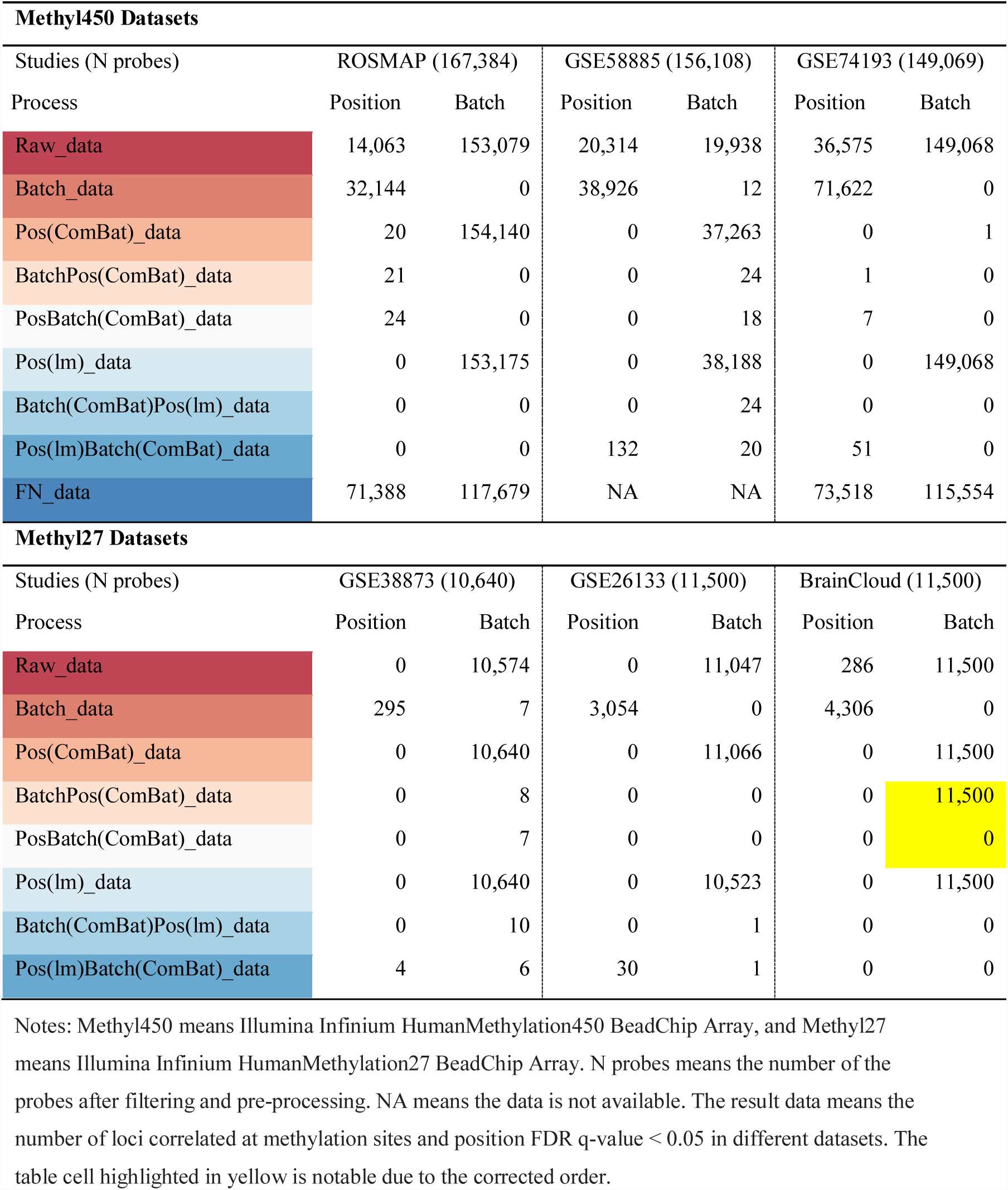
The number of CpG sites showed association with sample physical positions and batches in studies (FDR q-value<0.05). We used Analysis of Variance (ANOVA) analysis to calculate the p-values of correlation between methylation levels and position or batch. FDR q-value was computed for each nominal p-value by controlling the false discovery rate at 0.05 using the R function *qvalue.* We then obtained the number of CpGs significantly associated with positions and batches.

| **Methyl450 Datasets** |  |  |  |  |  |  |
| --- | --- | --- | --- | --- | --- | --- |
| Studies (N probes) | ROSMAP (167,384) |  | GSE58885 (156,108) |  | GSE74193 (149,069) |  |
| Process | Position | Batch | Position | Batch | Position | Batch |
| Raw_data | 14,063 | 153,079 | 20,314 | 19,938 | 36,575 | 149,068 |
| Batch_data | 32,144 | 0 | 38,926 | 12 | 71,622 | 0 |
| Pos(ComBat)_data | 20 | 154,140 | 0 | 37,263 | 0 | 1 |
| BatchPos(ComBat)_data | 21 | 0 | 0 | 24 | 1 | 0 |
| PosBatch(ComBat)_data | 24 | 0 | 0 | 18 | 7 | 0 |
| Pos(lm)_data | 0 | 153,175 | 0 | 38,188 | 0 | 149,068 |
| Batch(ComBat)Pos(lm)_data | 0 | 0 | 0 | 24 | 0 | 0 |
| Pos(lm)Batch(ComBat)_data | 0 | 0 | 132 | 20 | 51 | 0 |
| FN_data | 71,388 | 117,679 | NA | NA | 73,518 | 115,554 |
| **Methyl27 Datasets** |  |  |  |  |  |  |
| Studies (N probes) | GSE38873 (10,640) |  | GSE26133 (11,500) |  | BrainCloud (11,500) |  |
| Process | Position | Batch | Position | Batch | Position | Batch |
| Raw_data | 0 | 10,574 | 0 | 11,047 | 286 | 11,500 |
| Batch_data | 295 | 7 | 3,054 | 0 | 4,306 | 0 |
| Pos(ComBat)_data | 0 | 10,640 | 0 | 11,066 | 0 | 11,500 |
| BatchPos(ComBat)_data | 0 | 8 | 0 | 0 | 0 | 11,500 |
| PosBatch(ComBat)_data | 0 | 7 | 0 | 0 | 0 | 0 |
| Pos(lm)_data | 0 | 10,640 | 0 | 10,523 | 0 | 11,500 |
| Batch(ComBat)Pos(lm)_data | 0 | 10 | 0 | 1 | 0 | 0 |
| Pos(lm)Batch(ComBat)_data | 4 | 6 | 30 | 1 | 0 | 0 |

Notes: Methyl450 means Illumina Infinium HumanMethylation450 BeadChip Array, and Methyl27 means Illumina Infinium HumanMethylation27 BeadChip Array. N probes means the number of the probes after filtering and pre-processing. NA means the data is not available. The result data means the number of loci correlated at methylation sites and position FDR q-value < 0.05 in different datasets. The table cell highlighted in yellow is notable due to the corrected order.

We next attempted to correct the positional effects by the linear regression method^27, 35, 36^, through *lm* function in R. Regardless the orders of how the positional and batch effects were corrected, there were no CpG sites related to the physical positions. We normalized the data by functional normalization. In the ANOVA evaluated results, the FN_data did not perform well. Detailed results are shown in Table 2. We further analyzed another two Methyl450 and three Methyl27 datasets (See Materials and Methods, Table 3), and confirmed the existence of positional effects in those data (See Table 2).

**Table 3.** Information of verified datasets we used in this study. We used two Methyl450 datasets and three Methyl27 datasets from public databases and our own data to verify the results.

| Datasets<br>(NSAMPLES) | Methyl450 Datasets |  | Methyl27 Datasets |  |  |
| --- | --- | --- | --- | --- | --- |
|  | GSE58885<br>(179) | GSE74193<br>(673) | GSE38873<br>(153) | BrainCloud<br>(106) | GSE26133<br>(160) |
| **Tissue** | FC | DLPC (BA46/9) | CRBLM | CRBLM | LCLs |
| **Age, year** | -0.25± 0.07 | 36.13± 22.92 | 44.27± 9.84 | 35.86± 23.62 | NA |
| **Sex** |  |  |  |  |  |
| **Female** | 79 | 244 | 57 | 51 | 90 |
| **Male** | 100 | 429 | 96 | 55 | 70 |
| **Race** |  |  |  |  |  |
| **White** | NA | 317 | 153 | 42 | 0 |
| **AA** | NA | 356 | 0 | 64 | 0 |
| **African** | NA | 0 | 0 | 0 | Yoruba 160 |
| **Affection status** |  |  |  |  |  |
| **Control** | 179 | 224 | 47 | 106 | 160 |
| **Schiz** | 0 | 449 | 45 | 0 | 0 |
| **Dep** | 0 | 0 | 15 | 0 | 0 |
| **BP** | 0 | 0 | 46 | 0 | 0 |

Notes: NSAMPLES, # of samples; NA, Not available; FC, frontal cortex; DLPC, Dorsolateral prefrontal cortex; BA, Brodmann area; CRBLM, Cerebellum; Schiz, Schizophrenia; Dep, Depression; BP, Bipolar; AA, African American; LCLs, lymphoblastoid cell lines.

We further assessed the impact of the processes controlling batch and positional effects had on the data. We calculated the average methylation levels of ROSMAP data comparing pre- and post- correction in twelve positions and two batches respectively. After correcting the batches and positions by *ComBat* regardless of order, the methylation levels in the twelve physical positions became homogeneous (Fig.2a), and the differences of batch correction results remained statistically insignificant (Fig.2b). Alternatively, when we corrected positional effects by linear regression and functional normalization method, the variation of methylation levels in different physical positions had no significant reduction (Fig.2a); the same was seen for the batches (Fig.2b). The difference between sample locations was normalized after removing positional effects by *ComBat*. Similar results are also displayed in the Supplemental Materials (Fig.S2a, Fig.S2b).

**Figure 2.**
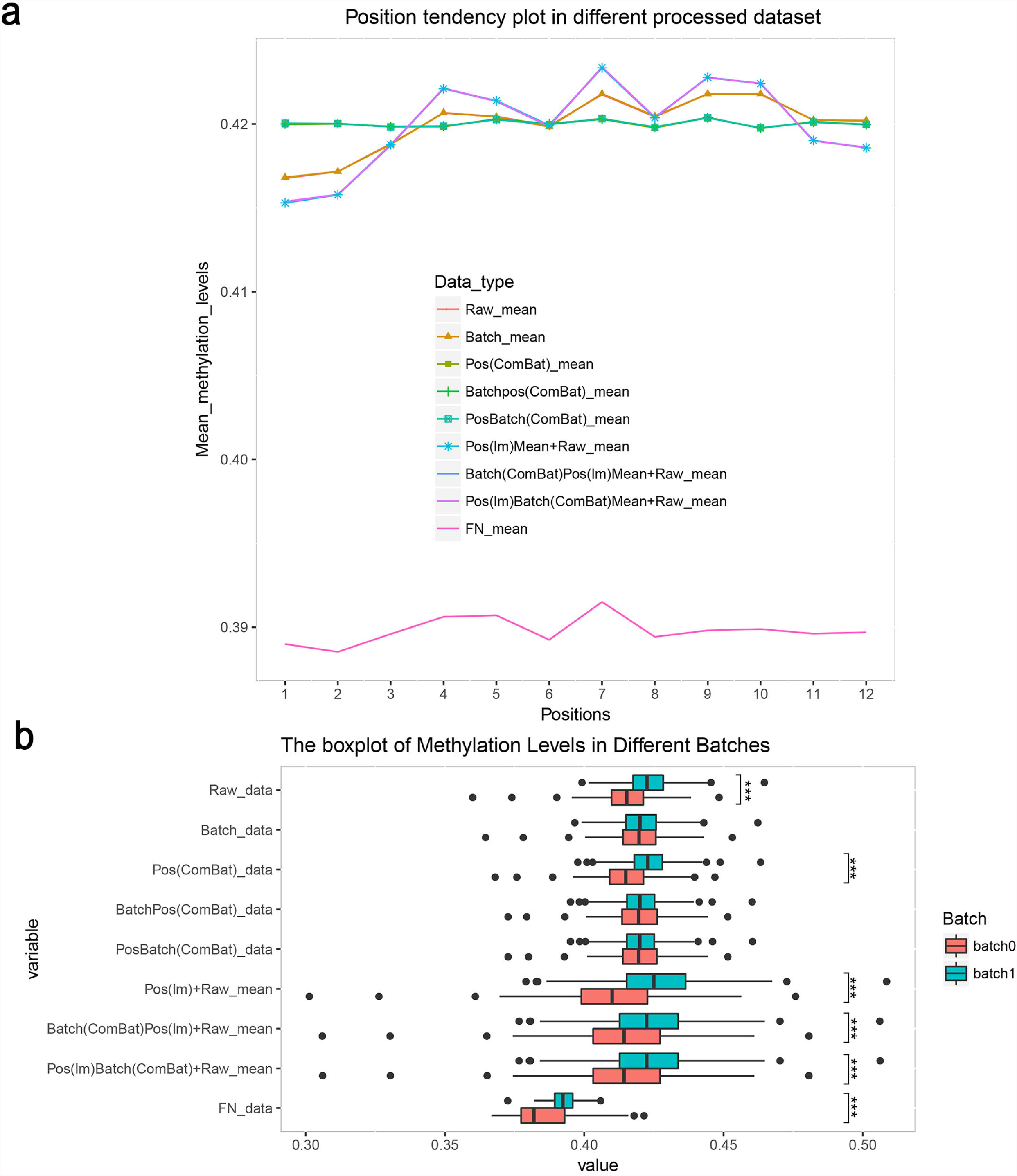
The average and variation of methylation levels of all probes in ROSMAP eight different processed datasets. (a) Average methylation levels in different positions. (b) Methylation levels in different batches. The corrected results by lm mean the residual of the linear regression model, so the raw mean is added to make the data in a normal range. The p-value was calculated using t-test and three stars mean the difference was statistically significant.

### Principal Variance Component Analysis (PVCA)

We made the PVCA plot to evaluate the relative weighted proportion variance (Fig.3). The PVCA plot describes the relative weights of corresponding eigenvectors related to the eigenvalues that can be explained by factors in the experimental design and other covariates^37, 38^. Here we considered eleven possible sources of variation: the two types of cell; age at cycle – baseline (age_bl) which can be the cognitive date, interview date, or clinical evaluation; age at death (age_death); the education level (educ); the cognitive diagnosis (cogdx); race (race); Spanish ancestry (spanish); sex; batch; and positional effects (position) and the weight of residual effect (resid in the figure) caused by unexplainable factors.

**Figure 3.**
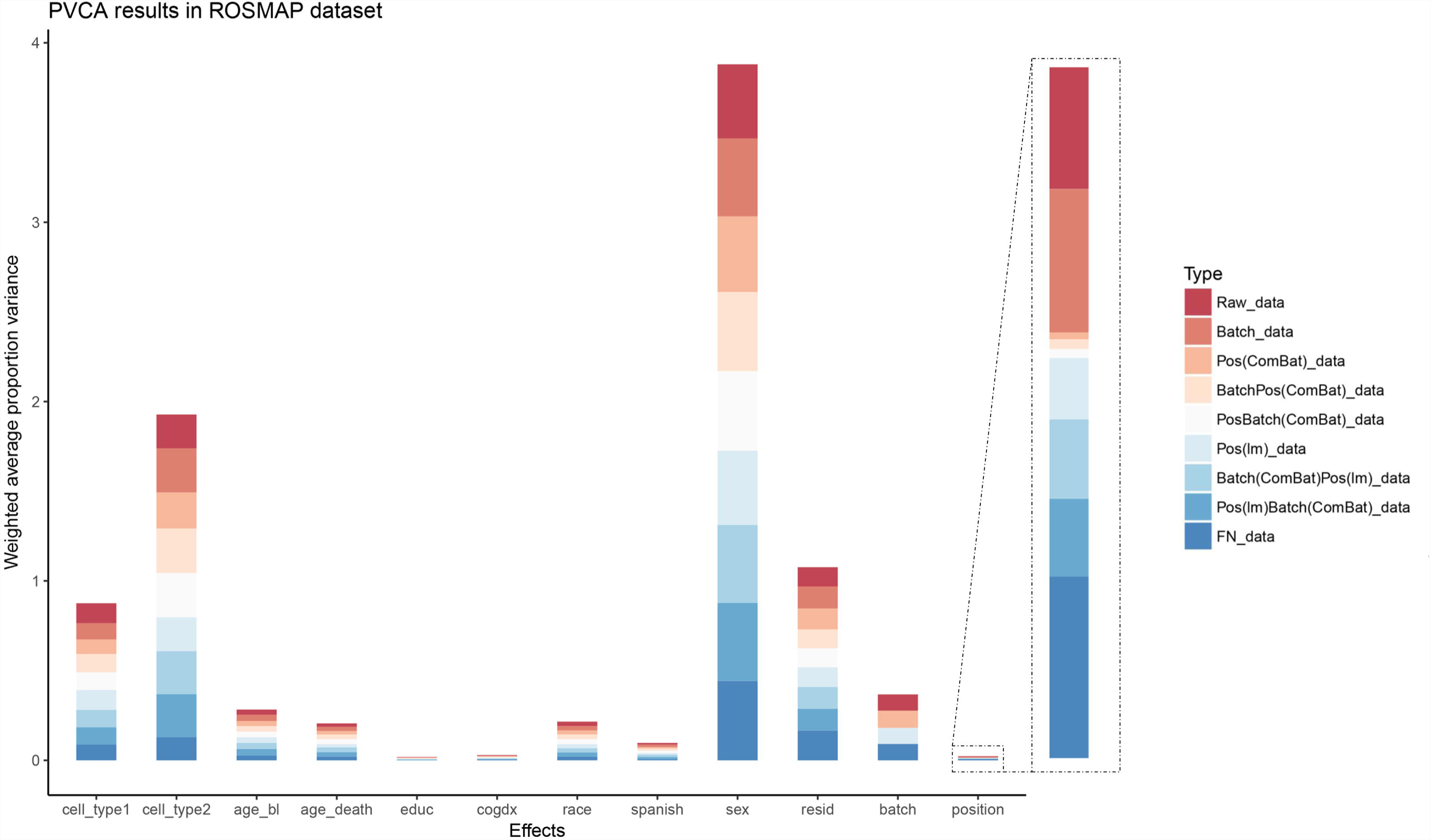
PVCA (principal variance component analysis) results in ROSMAP dataset. PVCA estimated the contribution of each factor to the overall variation. We considered eleven possible sources of variation in the ROSMAP data: the two types of cell; age at cycle – baseline (age_bl) which can be the cognitive date, interview date, or clinical evaluation; age at death (age_death); the education level (educ); the cognitive diagnosis (cogdx); race (race); Spanish ancestry (spanish); sex; batch; and positional effects (position). And we got the weight of residual effect (resid) that known factors could not explain.

The PVCA plot revealed that the BatchPos(ComBat)_data and PosBatch(ComBat)_data performed well in these nine datasets. Because these two datasets perform well in the technical variants both in batch effects and positional effects. By comparing the weighted proportional variance, we found the *ComBat* method outperformed *lm* in controlling the positional effects.

Nevertheless, other datasets reinforced the observation of positional effects. The similar results from the other datasets are displayed in the Supplemental Materials (Fig.S4, S5). These data indicate that the positional effects gave a relatively smaller contribution to the overall variation than other major factors like sex, age, and race, but it is not negligible.

### Analysis of the Technical Replicates

Technical replicates can be used to evaluate the consistency or precision of measurement. With this in mind, we considered whether removing positional effects can improve precision. The GSE74193 dataset had 140 pairs of technical replicates, and the GSE26133 dataset had 83 pairs. Subsequent to each correction step, the correlation values of each pair were calculated (Fig.4, and Fig.S6). After removing the positional effects by *ComBat*, the correlation increased more than lm (Wilcoxon signed-rank one-tailed test, p-value<2.2E-16) (Fig.4a, 4b, 4c and Fig.S6a, S6b, S6c) and functional normalization (Wilcoxon signed-rank one-tailed test, p-value = 0.0002) (Fig.S6d). Therefore, *ComBat* outperforms *lm* and functional normalization in adjusting the positional effects and improving precision. The correlation values in PosBatch(ComBat)_data is higher than BatchPos(ComBat)_data (Wilcoxon signed-rank one-tailed test, p-value=2.059E-15) (Fig.4d); thus correcting the positional effects first followed by batch effect improves precision. As for the efficiency of correction of positional effects, correcting the positional effects and batch effects could improve the correlation of technical replicates pairs (Wilcoxon signed-rank one-tailed test, p-value<2.2E-16) (Fig.S6g). However, there is no significant difference in correlations between data corrected for the positional effects before batch and data corrected for the batch only (Wilcoxon signed-rank one-tailed test, p-value=0.8667) (Fig.4e and Fig.S6f).

**Figure 4.**
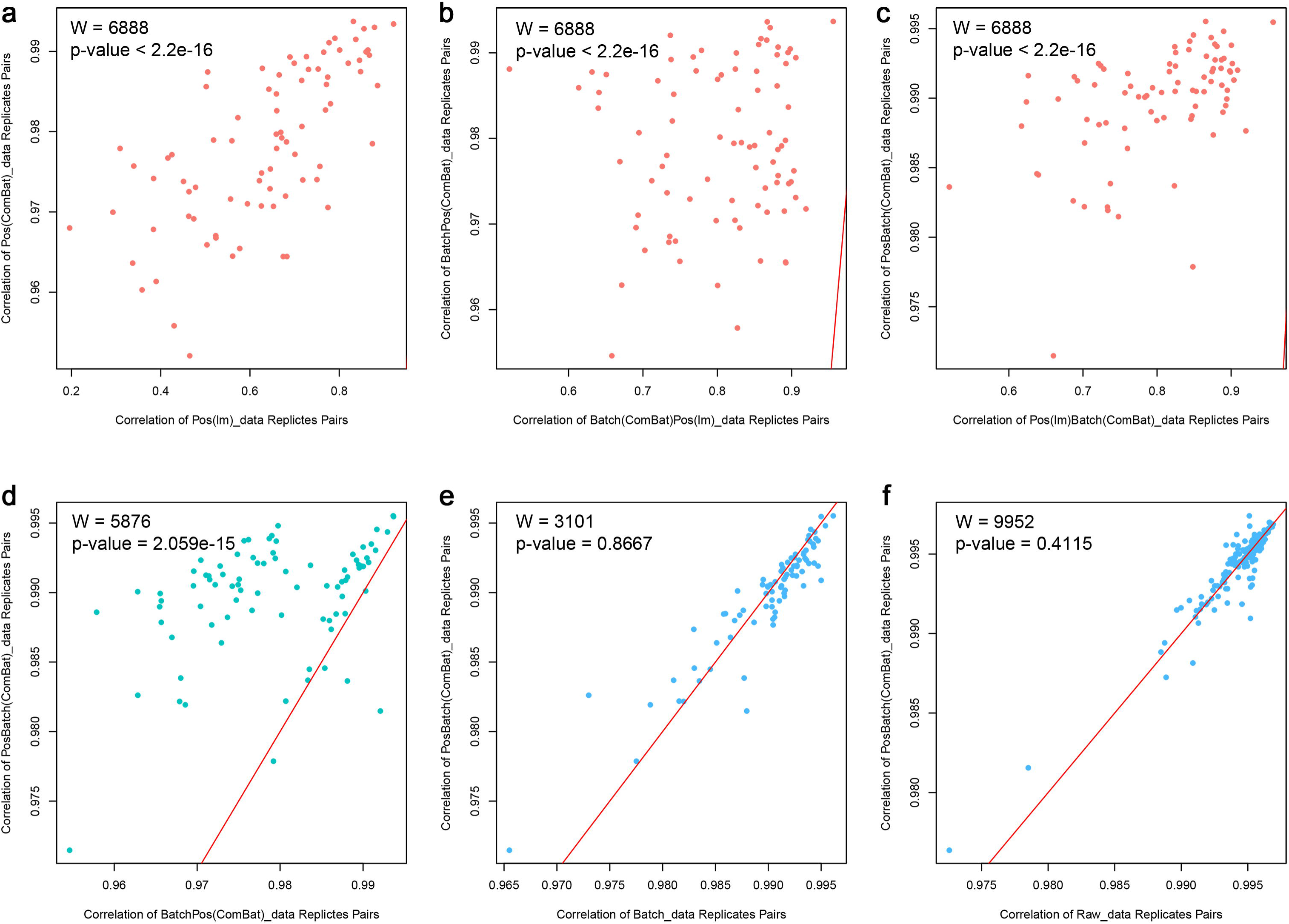
The comparison between different processed datasets in GSE26133. The Fig.4a, 4b and 4c are used to compare the correction methods of positional effects, the Fig.4d is used to compare the correction order of positional effects and batch effect, the Fig.4e and 4f figures are used to evaluate the efficiency of correction of positional effects. (a) Pos(ComBat)_data versus Pos(lm)_Data, (b) BatchPos(ComBat)_data versus Batch(ComBat)Pos(lm)_data, (c) PosBatch(ComBat)_data versus Pos(lm)Batch(ComBat)_data, (d) PosBatch(ComBat)_data versus BatchPos(ComBat)_data, (e) PosBatch(ComBat)_data versus Batch_data, (f) PosBatch(ComBat)_data versus Raw_data. The red lines mean y=x. The top left corner values reveal the Wilcoxon signed-rank one-tailed test result, the W is a test statistic means the sum of the signed ranks, which can be compared to a critical value from a reference table to get a p-value.

In summary, the best practice to correct the positional effect is *ComBat*. Correcting the positional effects first followed by batch effect is better than the reverse order.

### Differential Methylation CpG Loci Analysis

The impact of positional effects on the detection of differential methylation signals was assessed. An empirical Bayes test, *limma* in R, was used to identify differentially methylated CpGs between cases and controls of the processed GSE74193 dataset with 46 controls and 30 cases in two replicated groups; the number of CpGs associated with the schizophrenia was noted (p-value < 0.05 among all CpGs analyzed). The result is quantified using the area under a receiver operating characteristic curve (AUC of ROC), a popular measure of the accuracy. A higher AUC was identified in the PosBatch(ComBat)_data and BatchPos(ComBat)_data compared with other processed datasets (DeLong’s test for two ROC curves, Pos_data vs. Poslm_data p-value = 0.05, Pos_data vs. FN_data p-value = 0.0005, PosBatch(ComBat)_data vs. Pos_data p-value = 3.818e-16, PosBatch(ComBat)_data vs. BatchPos(ComBat)_data p-value = 0.9682).

The ROSMAP dataset was also be used to identify differentially methylated CpGs. 167,384 probes have been tested for differential methylation after filtering. 1,839 of the CpG loci were differentially methylated in data corrected for the batch effects (FDR <0.05). 1,846 CpG loci were significant in data corrected for positional effects followed by batch correction (Fig.5a). There are 145 CpG loci that were detected in the Batch_data, but not in the PosBatch(ComBat)_data, and 152 CpG loci detected in the PosBatch(ComBat)_data, but not the Batch_data. One of 152 CpG loci named cg24519157 is located in gene CASS4, which is an AD significant signal studied in several studies^39-44^. Therefore the positional effects could have confounded the methylation comparisons between case and control if the positional effects were not corrected during pre-processing, subsequently producing errors, including false negatives. When examining the sample plating, we noticed that the cases and controls had not been randomly placed in each position. Some positions have more cases than the others. Basically, position 4 and 5 have the largest proportion differences (Fig.5b). No matter how optimal the processing is, without proper randomization of an experiment the data may produce bias in the analysis ^24, 45^.

**Figure 5.**
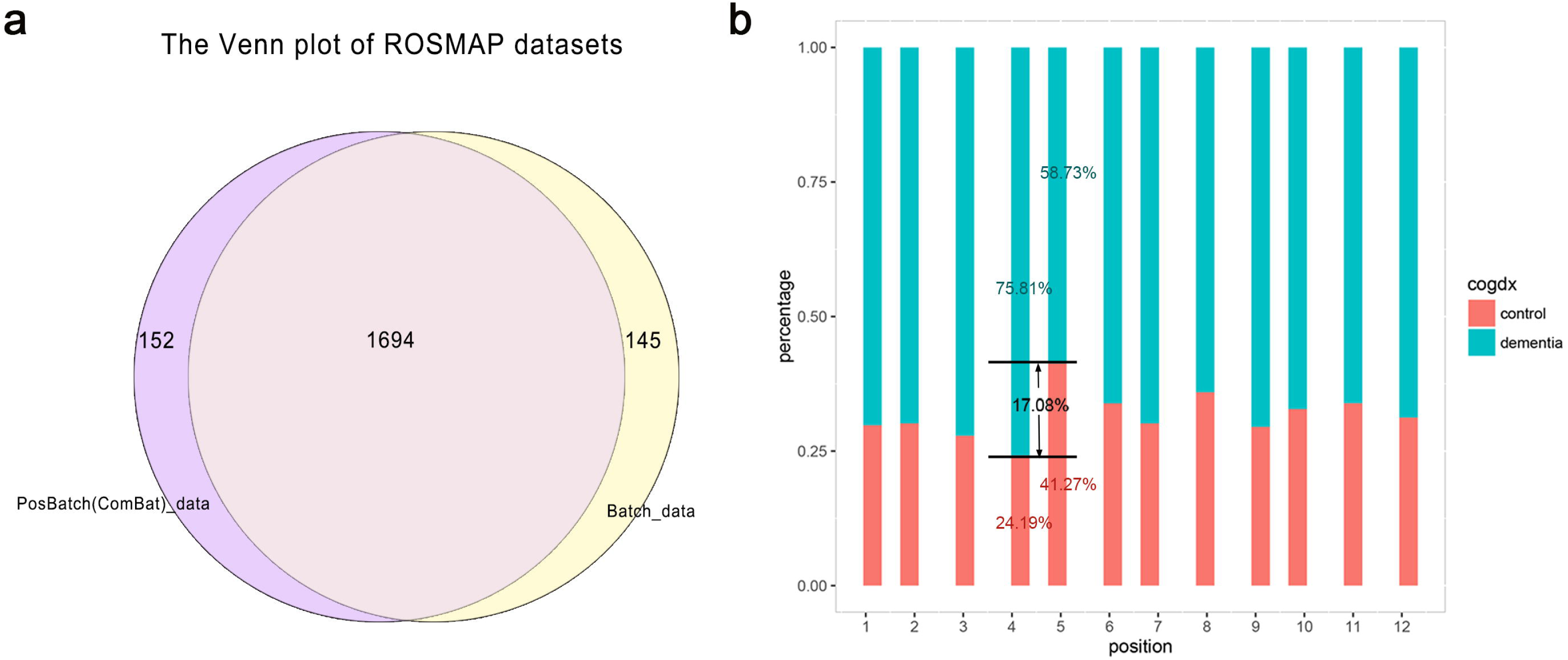
Differential Methylation CpG loci analysis results in ROSMAP dataset. (a) The Venn diagram plot of the differential methylated CpG loci obtained from differently processed data. The datasets including Batch_data (Data corrected for the batch effect), and PosBatch(ComBat)_data (Data corrected for the positional and batch effect sequentially by ComBat). (b) The sample distribution in cognitive diagnostic of ROSMAP data in twelve positions. The values showed in the figure is the control and dementia samples percentages in Position 4 and Position 5. Venn diagram showing the number of differential methylated CpGs (FDR< 0.05) between each pair of datasets.

## Discussion

Our analysis clearly identified an important technical artifact of the Illumina Infinium HumanMethylation BeadChips in both Methyl450 and Methyl27. We also studied one Illumina Infinium MethylationEPIC BeadChip microarray dataset (GEO: GSE86831) with 11 samples. 88,319 CpG loci of 864,935 loci were significantly associated with physical positions (FDR<0.05) in Batch_data, which suggest that the Illumina Infinium MethylationEPIC BeadChip microarray is also affected by the positional effects.

Due to the fact that positional effects can produce possible false conclusions, particular attention needs to be paid in controlling this variable in data analysis. In the analysis, we noticed the effect of positional effects in Methyl450 is larger than in Methyl27. The Type II probes contribute 2.98 times more to the positional effects than the Type I probes in Methyl450 datasets. After we had corrected the bias by Beta Mixture Quantile dilation (BMIQ, see *Material and methods*), a method used to adjust probe type bias, the proportion of Type II probes and Type I probes which were associated with position was 1.11. Adjusting the Type II probes methylation levels into a statistical distribution characteristic of Type I probes could help to reduce the positional effects, though the best method to remove the technical effects is correcting the positional effect by *ComBat* first and then removing the batch effect.

Although the technical replicates pairs could not prove the necessity of correcting the positional effects, the ANOVA results and differential CpGs analysis demonstrate that correcting the batch effect affect the positional effects and bias at many of the CpG loci. Therefore adjusting the positional effects is needed.

Two primary reasons for choosing the *ComBat* function in R to correct for the positional effect: first, the positional effect is randomly distributed in the same pattern as the batch effect, which *ComBat* is considered to be the most efficient method of correction^29-33^. Second, the correlation of technical replicate pairs and PVCA illustrates that the *ComBat* function is better than the linear regression model for correcting the positional effect. As for the correction order of positional and batch effects, we suggest correcting the positional effect first because of the first three evaluated methods.

In summary, positional effects can undoubtedly introduce bias into methylation level measures and produce unwanted bias. Sample placement in each chip should be randomized certainly^24, 45^, and most importantly, proper statistical methods should be used to remove the confounding artifacts. If the artifact is not taken into consideration, misleading conclusions could be drawn. Given that hundreds of epigenetics studies have used these platforms without controlling for positional effects, re-analysis of those previously published data and re-examining those reported significant signals may be needed.

## Material and methods

We have collected six datasets to test for positional effects. The datasets include three Methyl450 datasets and three Methyl27 datasets.

### Methyl450 Datasets

The primary data used in this study was a brain DNA collection obtained from Rush Alzheimer’s Disease Center in healthy controls and patients with dementia^46, 47^. The samples included 236 healthy controls and 507 dementia samples from two longitudinal cohort studies at Rush University Medical Center – the Religious Orders Study and the Rush Memory and Aging Project (ROSMAP data). The detailed sample information and the analysis pipeline were described in De Jager, P. L. et al., (2014) and Bennett, D. A. et al., (2012). The ROSMAP data was generated using the Methyl450 dataset and a sample of dorsolateral prefrontal cortex obtained from each sample. The sample information is listed in Table 1.

We used two other Methyl450 datasets to validate our findings: 179 frontal cortex samples from human fetal brains^11^ (GEO: GSE58885) (Genome Studio followed by WateRmelon in R. Normalized beta values generated via the Dasen method of the *WateRmelon* package); and 675 brain dorsolateral prefrontal cortex samples from Hernandez-Vargas’s study^48^ (GEO: GSE74193), which included 191 schizophrenia patients and 335 controls, with 140 technical replicates pairs or triplets included. Information on the two datasets is summarized in Table 3. And the ROSMAP and GSE74193 datasets have the .idat file, a binary format containing the raw red and green channel intensities.

### Methyl27 Datasets

We also used three Methyl27 datasets to confirm the findings. The datasets included the following: 153 cerebellum samples from our previous study (GEO: GSE38873)^49^, 106 brain prefrontal cortex samples from BrainCloud (downloaded from http://braincloud.jhmi.edu/downloads.htm)^50^, and 160 samples from GSE26133 with 83 technical replicates pairs, triplets or clusters included^51^. The beta-values of these studies were used directly to assess slide batch and positional effects. The information of these datasets was also summarized in Table 3.

### Data Quality Control and Pre-processing

We processed and analyzed data by R statistical language (www.r-project.org). The main processing pipeline is shown in Figure 1a (Fig. 1a). We removed probes and samples by detecting p-values obtained from GenomeStudio (Illumina, San Diego, CA). Samples were removed for those with more than 1% probes not detected (detection p-value > 0.01). We removed the probes with a bead count less than 3 in at least 5% of samples and probes with a detection p-value above 0.01 in more than one sample. At the end, 463,641 out of 485,577 probes remained for further analysis (Fig. 5a).

We then replaced the valued of 0 to 0.000001. Missing β value was imputed using a k-nearest neighbor algorithm by R *impute.knn* function in the *impute* package. To address the differences between the two types of probes, we used BMIQ (Beta Mixture Quantile dilation) function in *wateRmelon* package to adjust the β values of type II probes into a statistical distribution characteristic of type I probes, which has previously been shown to best minimize the variability between replicates^16, 52^.

The single nucleotide polymorphisms (SNPs) based on the 1000 Genomes database, small insertions and deletions (INDELs), repetitive DNA, and regions with reduced genomic complexity may affect the probe hybridization by a subject’s genotype^23^. After filtering, 167,384 probes remained for downstream analysis. The package *RefFreeEWAS* was utilized to estimate cell proportion^53^, and function *lm* was used to correct for the cellular heterogeneity.

The Methyl27 datasets were processed by the same pipeline as with Methyl450 datasets except the need to correct the probe type bias.

### Correct the Positional Effects

Here we used three methods to correct the positional effects: *ComBat* function, linear regression correction approach by using the *lm* function and the functional normalization (FN) method by using *preprocessFunnorm* function in *minfi* package (Fig.1b).

In our past studies, we found that the *ComBat* function in the R package *sva* is effective in removing the batch effects^29-33^. The function uses an empirical Bayesian method to adjust for known batches. Here we treated the positions as the batch information, and used the *ComBat* function applied to the high-dimensional data matrix, passing the full model matrix created without any known position variables. Position variables are passed as a separate argument to the function^54^, and the output is a set of corrected measurements where positional effects have been removed. We also used a linear regression model to adjust positions, and added the residues to the mean values as the corrected results. The FN method was also used to remove the positional effects^28^. It extends the idea of quantile normalization and uses control probes as surrogates. The method could be used to correct the positional effect and batch effect, mentioned in the Jean-Philippe Fortin. et al., (2014). It’s also worth noting that the functional normalization method can only be used for the 450K data with .idat files.

In addition, we attempted to modify the technique of calibrating variants like batch effects and positional effects (Fig. 1b). We built an R package to remove the positional effects and batch effects based on the *ComBat* function, named “posibatch”, to help you can correct these two confounders more easily. The package can be downloaded through https://github.com/ChuanJ/posibatch.

### Positional Effects Assessment

We used several metrics to evaluate positional effects for each dataset:

1. *The number of CpG loci significantly associated with positions.* We used Analysis of Variance analysis (ANOVA) to calculate the p-values of correlation between methylation levels and position or batch. FDR q-value was computed for each nominal p-value by controlling the false discovery rate at 0.05 using the R function *qvalue*^55^. We then obtained the number of CpGs significantly associated with positions and batches.
2. *A principal variance component analysis (PVCA) plot measured the attribution of impact factors to the methylation levels.* PVCA leverages the strengths of two statistic methods: principal components analysis (PCA) and variance components analysis (VCA). PCA is one of the most essential and popular techniques for reducing the dimensionality of a large dataset, increasing interpretability and minimizing information loss. VCA fits a linear mixed model to match the random effects to the factors of interest for estimating and partitioning the total variations. We made a PVCA plot by the *lme4* package in R to assess which processes are the most efficacious to correct positional effects.
3. *The correlation of technical replicated pairs.* We used the same evaluation metrics as Hailong Meng et al^56^ to determine the adequacy of eight datasets mentioned above separately.
4. *Differential Methylation CpG loci analysis.* To assess the impact of positional effects on analytical results, we discovered differentially methylated loci associated with schizophrenia in GSE74193 data and ROSMAP data using the *limma* package in R^57^. The GSE74193 dataset was divided into discovery and validation subgroups, with 30 patients’ samples and 46 controls in each subgroup. The *limma* is used to identify the differentially methylated loci and get the fold change (FC) between cases and controls. The area under a receiver operating characteristic curve (AUC of ROC) is generally used to measure the accuracy. The curve is created by plotting the true positive rate and false positive rate at various threshold settings. We identified AUC for the prediction of high and low fold changes. The cut-off is p-value lower than 0.05 and the log(FC) higher than 0.02. The AUC of ROC was used to measure the internal consistency in each normalization method. The Delong’s test was used to compare the AUC of ROC Curves^58^.

## Acknowledgements

We sincerely thank Chicago Biomedical Consortium for its supports (to C. Liu) as well. We are grateful to Zhanlin Chen, Yi Jiang and Lingling Huang for editing the manuscript. All the data contributors are also sincerely thanked for data submitted in the GEO, particularly Dr. Gregory Klein and David A Bennett for sharing their data of Religious Orders Study and Memory and Aging Project (ROSMAP).

## Funding

This work was supported by the National Institutes of Health under Grant numbers 1 U01 MH103340-01, 1R01ES024988 to C. Liu; National Natural Science Foundation of China under Grant numbers 81401114, 31571312 to C. Chen.

## References

1. Jaenisch R, Bird A. Epigenetic regulation of gene expression: how the genome integrates intrinsic and environmental signals. Nat Genet 2003; 33 Suppl:245-54.

2. Jones PA. Functions of DNA methylation: islands, start sites, gene bodies and beyond. Nat Rev Genet 2012; 13:484-92.

3. Girardot M, Feil R, Lleres D. Epigenetic deregulation of genomic imprinting in humans: causal mechanisms and clinical implications. Epigenomics 2013; 5:715-28.

4. McCabe DC, Caudill MA. DNA methylation, genomic silencing, and links to nutrition and cancer. Nutr Rev 2005; 63:183-95.

5. Heyn H, Vidal E, Ferreira HJ, Vizoso M, Sayols S, Gomez A, Moran S, Boque-Sastre R, Guil S, Martinez-Cardus A, et al. Epigenomic analysis detects aberrant super-enhancer DNA methylation in human cancer. Genome Biol 2016; 17:11.

6. Boerno ST, Grimm C, Lehrach H, Schweiger MR. Next-generation sequencing technologies for DNA methylation analyses in cancer genomics. Epigenomics 2010; 2:199-207.

7. Robertson KD. DNA methylation and human disease. Nat Rev Genet 2005; 6:597-610.

8. Pogribny IP, Beland FA. DNA hypomethylation in the origin and pathogenesis of human diseases. Cell Mol Life Sci 2009; 66:2249-61.

9. Wilson AS, Power BE, Molloy PL. DNA hypomethylation and human diseases. Biochim Biophys Acta 2007; 1775:138-62.

10. Heyn H, Li N, Ferreira HJ, Moran S, Pisano DG, Gomez A, Diez J, Sanchez-Mut JV, Setien F, Carmona FJ, et al. Distinct DNA methylomes of newborns and centenarians. Proc Natl Acad Sci U S A 2012; 109:10522-7.

11. Spiers H, Hannon E, Schalkwyk LC, Smith R, Wong CC, O’Donovan MC, Bray NJ, Mill J. Methylomic trajectories across human fetal brain development. Genome Res 2015; 25:338-52.

12. Lim AS, Srivastava GP, Yu L, Chibnik LB, Xu J, Buchman AS, Schneider JA, Myers AJ, Bennett DA, De Jager PL. 24-hour rhythms of DNA methylation and their relation with rhythms of RNA expression in the human dorsolateral prefrontal cortex. PLoS Genet 2014; 10:e1004792.

13. Muangsub T, Samsuwan J, Tongyoo P, Kitkumthorn N, Mutirangura A. Analysis of methylation microarray for tissue specific detection. Gene 2014; 553:31-41.

14. Shanmuganathan R, Basheer NB, Amirthalingam L, Muthukumar H, Kaliaperumal R, Shanmugam K. Conventional and nanotechniques for DNA methylation profiling. J Mol Diagn 2013; 15:17-26.

15. Bibikova M, Le J, Barnes B, Saedinia-Melnyk S, Zhou L, Shen R, Gunderson KL. Genome-wide DNA methylation profiling using Infinium(R) assay. Epigenomics 2009; 1:177-200.

16. Dedeurwaerder S, Defrance M, Calonne E, Denis H, Sotiriou C, Fuks F. Evaluation of the Infinium Methylation 450K technology. Epigenomics 2011; 3:771-84.

17. Dedeurwaerder S, Defrance M, Bizet M, Calonne E, Bontempi G, Fuks F. A comprehensive overview of Infinium HumanMethylation450 data processing. Brief Bioinform 2014; 15:929-41.

18. Moran S, Arribas C, Esteller M. Validation of a DNA methylation microarray for 850,000 CpG sites of the human genome enriched in enhancer sequences. Epigenomics 2016; 8:389-99.

19. Bibikova M. High-throughput DNA methylation profiling using universal bead arrays. Genome Research 2006; 16:383-93.

20. Teh AL, Pan H, Lin X, Lim YI, Patro CP, Cheong CY, Gong M, MacIsaac JL, Kwoh CK, Meaney MJ, et al. Comparison of Methyl-capture Sequencing vs. Infinium 450K methylation array for methylome analysis in clinical samples. Epigenetics 2016; 11:36-48.

21. Bibikova M, Barnes B, Tsan C, Ho V, Klotzle B, Le JM, Delano D, Zhang L, Schroth GP, Gunderson KL, et al. High density DNA methylation array with single CpG site resolution. Genomics 2011; 98:288-95.

22. Chen YA, Lemire M, Choufani S, Butcher DT, Grafodatskaya D, Zanke BW, Gallinger S, Hudson TJ, Weksberg R. Discovery of cross-reactive probes and polymorphic CpGs in the Illumina Infinium HumanMethylation450 microarray. Epigenetics 2013; 8:203-9.

23. Naeem H, Wong NC, Chatterton Z, Hong MK, Pedersen JS, Corcoran NM, Hovens CM, Macintyre G. Reducing the risk of false discovery enabling identification of biologically significant genome-wide methylation status using the HumanMethylation450 array. BMC Genomics 2014; 15:51.

24. Verdugo RA, Deschepper CF, Munoz G, Pomp D, Churchill GA. Importance of randomization in microarray experimental designs with Illumina platforms. Nucleic Acids Res 2009; 37:5610-8.

25. Teschendorff AE, Zhuang J, Widschwendter M. Independent surrogate variable analysis to deconvolve confounding factors in large-scale microarray profiling studies. Bioinformatics 2011; 27:1496-505.

26. Moran S, Vizoso M, Martinez-Cardus A, Gomez A, Matias-Guiu X, Chiavenna SM, Fernandez AG, Esteller M. Validation of DNA methylation profiling in formalin-fixed paraffin-embedded samples using the Infinium HumanMethylation450 Microarray. Epigenetics 2014; 9:829-33.

27. Kulkarni H, Kos MZ, Neary J, Dyer TD, Kent JW, Jr., Goring HH, Cole SA, Comuzzie AG, Almasy L, Mahaney MC, et al. Novel epigenetic determinants of type 2 diabetes in Mexican-American families. Hum Mol Genet 2015; 24:5330-44.

28. Fortin JP, Labbe A, Lemire M, Zanke BW, Hudson TJ, Fertig EJ, Greenwood CM, Hansen KD. Functional normalization of 450k methylation array data improves replication in large cancer studies. Genome Biol 2014; 15:503.

29. Johnson WE, Li C, Rabinovic A. Adjusting batch effects in microarray expression data using empirical Bayes methods. Biostatistics 2007; 8:118-27.

30. Sun Z, Chai HS, Wu Y, White WM, Donkena KV, Klein CJ, Garovic VD, Therneau TM, Kocher JP. Batch effect correction for genome-wide methylation data with Illumina Infinium platform. BMC Med Genomics 2011; 4:84.

31. Chen C, Grennan K, Badner J, Zhang D, Gershon E, Jin L, Liu C. Removing batch effects in analysis of expression microarray data: an evaluation of six batch adjustment methods. PLoS One 2011; 6:e17238.

32. Leek JT, Scharpf RB, Bravo HC, Simcha D, Langmead B, Johnson WE, Geman D, Baggerly K, Irizarry RA. Tackling the widespread and critical impact of batch effects in high-throughput data. Nat Rev Genet 2010; 11:733-9.

33. Lazar C, Meganck S, Taminau J, Steenhoff D, Coletta A, Molter C, Weiss-Solis DY, Duque R, Bersini H, Nowe A. Batch effect removal methods for microarray gene expression data integration: a survey. Brief Bioinform 2013; 14:469-90.

34. Barrett T, Troup DB, Wilhite SE, Ledoux P, Rudnev D, Evangelista C, Kim IF, Soboleva A, Tomashevsky M, Marshall KA, et al. NCBI GEO: archive for high-throughput functional genomic data. Nucleic Acids Res 2009; 37:D885-90.

35. Agha G, Houseman EA, Kelsey KT, Eaton CB, Buka SL, Loucks EB. Adiposity is associated with DNA methylation profile in adipose tissue. Int J Epidemiol 2015; 44:1277-87.

36. Marabita F, Almgren M, Lindholm ME, Ruhrmann S, Fagerstrom-Billai F, Jagodic M, Sundberg CJ, Ekstrom TJ, Teschendorff AE, Tegner J, et al. An evaluation of analysis pipelines for DNA methylation profiling using the Illumina HumanMethylation450 BeadChip platform. Epigenetics 2013; 8:333-46.

37. Harrison JM, Howard D, Malven M, Halls SC, Culler AH, Harrigan GG, Wolfinger RD. Principal variance component analysis of crop composition data: a case study on herbicide-tolerant cotton. Journal of agricultural and food chemistry 2013; 61:6412-22.

38. Boedigheimer MJ, Wolfinger RD, Bass MB, Bushel PR, Chou JW, Cooper M, Corton JC, Fostel J, Hester S, Lee JS, et al. Sources of variation in baseline gene expression levels from toxicogenomics study control animals across multiple laboratories. BMC Genomics 2008; 9:285.

39. Cruts M, Theuns J, Van Broeckhoven C. Locus-specific mutation databases for neurodegenerative brain diseases. Hum Mutat 2012; 33:1340-4.

40. Deneka A, Korobeynikov V, Golemis EA. Embryonal Fyn-associated substrate (EFS) and CASS4: The lesser-known CAS protein family members. Gene 2015; 570:25-35.

41. Beck TN, Nicolas E, Kopp MC, Golemis EA. Adaptors for disorders of the brain? The cancer signaling proteins NEDD9, CASS4, and PTK2B in Alzheimer’s disease. Oncoscience 2014; 1:486-503.

42. Karch CM, Goate AM. Alzheimer’s disease risk genes and mechanisms of disease pathogenesis. Biol Psychiatry 2015; 77:43-51.

43. Chouraki V, Seshadri S. Genetics of Alzheimer’s disease. Adv Genet 2014; 87:245-94.

44. Wang X, Lopez OL, Sweet RA, Becker JT, DeKosky ST, Barmada MM, Demirci FY, Kamboh MI. Genetic determinants of disease progression in Alzheimer’s disease. J Alzheimers Dis 2015; 43:649-55.

45. Triche TJ, Jr., Weisenberger DJ, Van Den Berg D, Laird PW, Siegmund KD. Low-level processing of Illumina Infinium DNA Methylation BeadArrays. Nucleic Acids Res 2013; 41:e90.

46. De Jager PL, Srivastava G, Lunnon K, Burgess J, Schalkwyk LC, Yu L, Eaton ML, Keenan BT, Ernst J, McCabe C, et al. Alzheimer’s disease: early alterations in brain DNA methylation at ANK1, BIN1, RHBDF2 and other loci. Nat Neurosci 2014; 17:1156-63.

47. Bennett DA, Schneider JA, Arvanitakis Z, Wilson RS. Overview and findings from the religious orders study. Current Alzheimer research 2012; 9:628-45.

48. Jaffe AE, Gao Y, Deep-Soboslay A, Tao R, Hyde TM, Weinberger DR, Kleinman JE. Mapping DNA methylation across development, genotype and schizophrenia in the human frontal cortex. Nat Neurosci 2016; 19:40-7.

49. Zhang D, Cheng L, Badner JA, Chen C, Chen Q, Luo W, Craig DW, Redman M, Gershon ES, Liu C. Genetic control of individual differences in gene-specific methylation in human brain. Am J Hum Genet 2010; 86:411-9.

50. Numata S, Ye T, Hyde TM, Guitart-Navarro X, Tao R, Wininger M, Colantuoni C, Weinberger DR, Kleinman JE, Lipska BK. DNA methylation signatures in development and aging of the human prefrontal cortex. Am J Hum Genet 2012; 90:260-72.

51. Bell JT, Pai AA, Pickrell JK, Gaffney DJ, Pique-Regi R, Degner JF, Gilad Y, Pritchard JK. DNA methylation patterns associate with genetic and gene expression variation in HapMap cell lines. Genome Biol 2011; 12:R10.

52. Teschendorff AE, Marabita F, Lechner M, Bartlett T, Tegner J, Gomez-Cabrero D, Beck S. A beta-mixture quantile normalization method for correcting probe design bias in Illumina Infinium 450 k DNA methylation data. Bioinformatics 2013; 29:189-96.

53. Houseman EA, Kelsey KT, Wiencke JK, Marsit CJ. Cell-composition effects in the analysis of DNA methylation array data: a mathematical perspective. BMC Bioinformatics 2015; 16:95.

54. Leek JT, Johnson WE, Parker HS, Jaffe AE, Storey JD. The sva package for removing batch effects and other unwanted variation in high-throughput experiments. Bioinformatics 2012; 28:882-3.

55. Storey JD, Tibshirani R. Statistical significance for genomewide studies. Proc Natl Acad Sci U S A 2003; 100:9440-5.

56. Meng H, Joyce AR, Adkins DE, Basu P, Jia Y, Li G, Sengupta TK, Zedler BK, Murrelle EL, van den Oord EJ. A statistical method for excluding non-variable CpG sites in high-throughput DNA methylation profiling. BMC Bioinformatics 2010; 11:227.

57. Ritchie ME, Phipson B, Wu D, Hu Y, Law CW, Shi W, Smyth GK. limma powers differential expression analyses for RNA-sequencing and microarray studies. Nucleic Acids Res 2015; 43:e47.

58. DeLong ER, DeLong DM, Clarke-Pearson DL. Comparing the areas under two or more correlated receiver operating characteristic curves: a nonparametric approach. Biometrics 1988; 44:837-45.

